# Deep learning: step forward to high-resolution in vivo shortwave infrared imaging

**DOI:** 10.1101/2021.03.04.433844

**Authors:** Vladimir A. Baulin, Yves Usson, Xavier Le Guével

## Abstract

Shortwave infrared window (SWIR: 1000-1700 nm) represents a major improvement compared to the NIR-I region (700-900 nm) in terms of temporal and spatial resolutions in depths down to 4 mm. SWIR is a fast and cheap alternative to more precise methods such as X-ray and opto-acoustic imaging. Main obstacles in SWIR imaging are the noise and scattering from tissues and skin that reduce the precision of the method. We demonstrate that the combination of SWIR in vivo imaging in the NIR-IIb region (1500-1700 nm) with advanced deep learning image analysis allows to overcome these obstacles and making a large step forward to high resolution imaging: it allows to precisely segment vessels from tissues and noise, provides morphological structure of the vessels network, with learned pseudo-3D shape, their relative position, dynamic information of blood vascularization in depth in small animals and distinguish the vessels types: artieries and veins. For demonstration we use neural network IterNet that exploits structural redundancy of the blood vessels, which provides a useful analysis tool for raw SWIR images.

## 1 INTRODUCTION

The field of *in vivo* optical imaging for biomedical applications is expanding rapidly over the last two decades leading to more precise diagnostic of early stage diseases and to advanced image-guided-surgery system already available for clinical applications^[1]^. One of these breakthroughs is related to the development of innovative imaging systems in the shortwave infrared (SWIR) spectral window, called also NIR-II, between 900 and 1700 nm. SWIR has demonstrated a major improvement in terms of spatial and temporal resolution, reaching deep in tissue down to 4 to 6 mm compared to the Visible (400-700 nm) and NIR-I (700-900 nm) regions. The benefit moving forward from NIR-I to SWIR is mainly associated to the weak auto-fluorescence and reduced scattering from the living tissues at longer wavelengths^[2]^. For instance, it was shown recently the striking improvement of detection with higher signal-to-noise ratio selecting the SWIR sub-windows NIR-IIb (1500-1700 nm) for *in vivo* imaging^[3–5]^. The concomitant progress of the sensor technology in the SWIR range and of the formulation of new bright and biocompatible SWIR emitting organic and inorganic contrast agents^[6–9]^ has enabled to use these optical systems for intra-operative surgery in small animals^[10,11]^ and recently in human^[12]^. One of the most appealing field of applications for SWIR imaging concerns the (micro)vascularization, with success to monitor in real time non-invasively different pathologies such as vascular disorders, (neo)angegiogenosis in cancer, wound healing, implants[7,13–15].

Despite these major steps, we are still far to reach the spatial resolution below one micron at high depth achieved by X-ray imaging^[16]^. Other recent optical imaging systems based on full field optical coherence^[17]^ and high-resolution optoacoustic imaging^[18]^ lead to spatial resolution down to 1.7 um but with a quite limit field of view that requires long time acquisition to image the whole animal. A promising strategy to overcome this issue relies on the image analysis using deep neural networks. This field shows exponential growth in recent years due to growth in computational power of modern parallel computers, and the quality of feature extraction, detection and segmentation made by deep neural networks has made a major step forward^[19–21]^. In particular, several networks built on a popular fully connected convolutional neural network (CNN) U-Net^[22]^, were very successfully applied in the context of a classical problem of the segmentation of retina vessels using as a training set a series of open annotated databases for retina vessels. As a result, several developed deep neural networks from the leaderboard show the performance above 97% precision tested against ground truth. Thus, a logical extension of these networks would be the application of these developments for SWIR images. Although the nature of the images is different: shadows of the visible light passing through the tissue versus fluorescent NIR signal directly from the vessels, the structure of the vessels representing continuous interconnected lines with a certain redundancy enables the generalization of the developed networks to the case of SWIR images. It allows segment the vessels from the background, reduce scattering light originated from the tissues, and detect 3D blood vessels structures, thus providing essential information for a full structural analysis via skeletonization of the vessel network and enhanced statistics: number of branching points, average length of the vessels, the thickness of the vessels, relative length of vessels of different categories, etc. Thus, combination of fast and relatively cheap SWIR method with advanced deep learning image analysis may contribute to fill the gap in resolutions between SWIR and X-ray.

Recent study has demonstrated the significant improvement of contrast and spatial resolution of SWIR in mice using Monte Carlo Restoration which enabled to perform segmentation analysis of small animal presenting vascular disorder^[23]^. In this work we explore the advantages of using deep neural networks specially designed to extract blood vessels structure trained on large datasets of retina vessels in order to predict the structure of vessels in SWIR images. We use one of the best neural networks in prediction of vessels structures IterNet^[24]^ on *in vivo* SWIR NIR-IIb imaging to demonstrate the high potential of this method to go one-step further to high-resolution optical imaging that could be easily transferred in clinics and hospitals.

## 2 DEEP NEURAL NETWORK FOR SWIR IMAGES

Fully connected convolutional neural network is a popular neural network U-Net^[22]^ using strong data augmentation to significantly reduce the number of training images. Its successor, deep neural network IterNet^[24]^ is build on U-Net and combines iteratively *N* -1 mini-networks U-Net after one segmentation with U-Net. It goes further in precision and uses the structural redundancy or self-similarity of blood vessels that allows the network to find obscured details of the vessel from the segmented vessel image itself, rather than from the raw input image. In fact, IterNet can learn from as few as 10–20 annotated images (ground truth) to provide a good accuracy.

IterNet is one of the leaders in leaderboard in ratings^[25]^ for the performance in segmentation of retina blood vessels. The performance is tested on open databases of blood retina vessels on three mainstream data sets, DRIVE^[26]^, CHASE_DB1 ^[27]^ and STARE^[28]^, which are used as as a gold standard for the performance benchmark and comparison between segmentation networks for blood vessels. It has a high accuracy measured in terms of the receiver operating characteristic curve (ROC), which is plotting True Positive Rate (TPR) versus False Positive Rate (FPR). This measure is implemented in TensorFlow^[29]^ and the corresponding Area Under the ROC Curve (AUC) gives a numerical measure of the performance of the network training. The provided training weights^[24]^ of IterNet network give AUCs of 0.9816, 0.9851, and 0.9881 for the classical benchmarks of the data sets of retina vessels: DRIVE, CHASE_DB1, STARE, respectively.

To test the performance of the network for *in vivo* SWIR images of nude mice, we have performed several setups with different cameras, lenses and different distances from the sample. The quality of prediction was then tested on *ex vivo* post-mortem sample with removed skin. This allows to evaluate the structure of vessels with high magnification camera in order to confirm the prediction of microvessels.

### 2.1 SWIR *in vivo* images

SWIR imaging was performed using a Princeton camera 640ST (900-1700 nm) coupled with a laser excitation source at *λ*= 808 nm (100 mW/cm^2^). We used a short-pass excitation filter at 1000 nm (Thorlabs) and a long-pass filter on the SWIR camera from Thorlabs (LP1500 nm). 25 mm or 50 mm lenses with numerical aperture (n.a) = 1.4 (Navitar) were used to focus on the mice placed at 30 cm working distance. 25 mm and 50 mm lenses provide a theoretical spatial resolution of 400 microns and 150 microns respectively. NMRI nude mice (Janvier, France) were anesthetized (air/isofluorane 4% for induction and 1.5% thereafter) and were injected intravenously via the tail vein (200 *µ*L of Indocyanine Green (ICG) at 500 *µ*M in PBS). *In vivo* SWIR imaging was performed using 25 mm or 50 mm lenses and LP1500 nm at different exposure times from 100 ms to 1 s.

### 2.2 Training of the neural network

We use IterNet^[24]^ for prediction of vessels in SWIR images (i) with the released by the authors universal pre-trained weights trained across multiple datasets^[24]^. Each database contains 40 images of retina vessels taken with optical camera. Human experts manually segmented the vessels in each image under guidance of trained and experienced medical doctor (ground truth)^[26]^. The criteria for marking the vessel was 70% confidence that the structure is the vessel. These images were divided into two equal datasets: one dataset was using for training of the network and another dataset was used for evaluation of the performance. The evaluation of the performance of the training was done using standard method in Tensorflow for binary classifiers based on receiver operating characteristic (ROC) curve, which is a plot of true positive vs. false positive rates: higher area under the curve corresponds to better performance. The original weights were used without additional fine-tuning; (ii) weights from specially trained network on manually annotated SWIR images used as a ground truth for detection, example is Fig. 1. These images were added to the training set of DRIVE database, which were gray scaled. The manual annotation of SWIR training examples was not complete and it was done for the test of robustness of the prediction of the network. The training was performed with the following parameters: batch_size=32,repeat=10, minimum_kernel=32, epochs=200, iteration=3, crop_size=128, stride_size=3 and the resulting performance is shown in Fig. 1 C): An example of the original image, an example of the annotated image and the resulting ROC curve showing the performance on SWIR images. Although the annotated images were not as elaborated as in the vessel databases and despite their small number, the network predictions have 0.9 AUC score on images and manually annotations unseen during training. This result demonstrates that the network reliably predicts the vessels at least comparable with the human annotation results on SWIR images.

**FIGURE 1.**
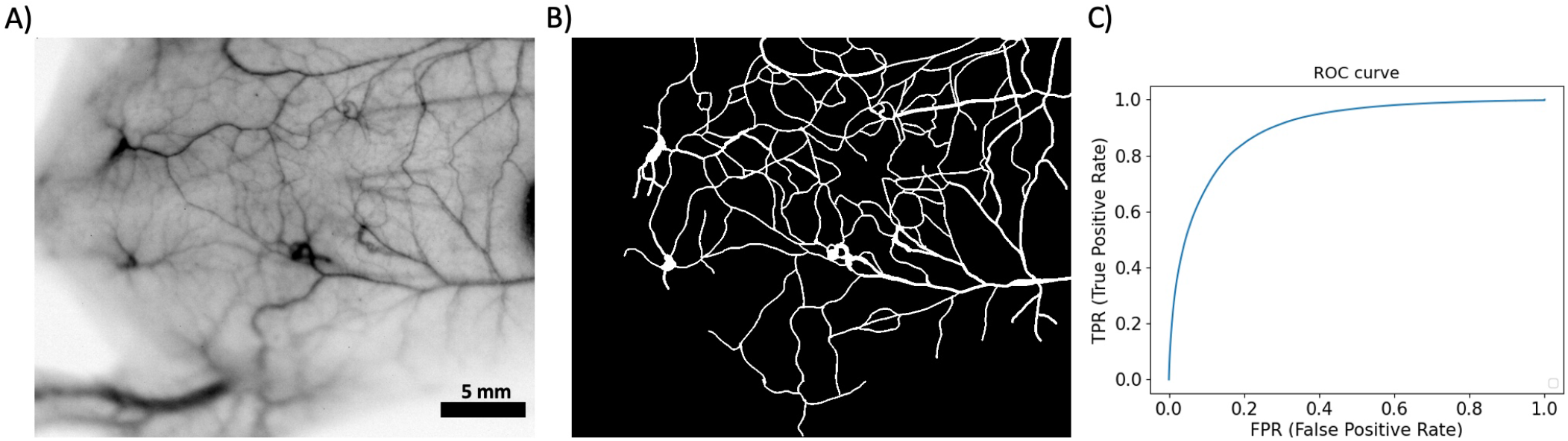
A) Original SWIR image used for training of the network and obtained with a 50 mm lens (n.a.= 1.4) at 100 ms exposure; B) Example of the annotation of ground truth, microvessels are not annotated; C) Result of the training in form of the receiver of operating characteristic (ROC) curve showing the performance of the training on SWIR images. Area under the curve: 0.90; Area under precision-recall curve: 0.57, Jaccard similarity: 0.89.

SWIR images have 640×512 pixels dimensions and the vessels are smaller than in the original training database. In order to achieve a good accuracy, SWIR images (dimensions of vessels and brightness/contrast) should be as close as possible to the training set used for the network training. SWIR images contain a large number of micro-vessels, almost invisible by eye. They were not annotated in the training set and in order to segment them the stride size and the crop size should be reduced to minimum. We use the following parameters that give the best results for the inference of the vessels with pre-trained weights: Activation=‘ReLU’, dropout=0.1, minimum_kernel=32, batch_size=128, epochs=600, iteration=3, stride_size=1, crop_size=16. The images were processed on a desktop computer equipped with AMD Ryzen 5 5600X processor with 64 GB RAM, Nvidia RTX 3090 graphical card with 24 GB memory, Ubuntu 20.04 and Tensorflow 1.15 ^[29]^ through Nvidia-TensorFlow Horovod^[30]^ compilation to ensure back compatibility with RTX 3090 graphical card. With this setup, the speed of processing of a SWIR image with all GPU overheads, pre-processing, post-processing and writing the resulting files to the disk is presented in Table 1.

**TABLE 1.** Approximate processing time of a SWIR image using Nvidia RTX 3090 graphical card. Optimization of the workflow would allow to make devices able to perform in near-real time vessels segmentation and analysis.

| Image resolution | Time of execution |
| --- | --- |
| $1280 \times 1024$ | 4 min 40 s |
| $640 \times 512$ | 1 min 10 s |
| $320 \times 256$ | 0 min 18 s |

### 2.3 Neural network prediction

The performance and the number of vessels predicted by the network largely depends on the quality of SWIR images, the distance of the camera from the mouse skin and the brightness/noise of the vessels. The example of the predicted vessels is shown in Fig. 2. The network can not only segment vessels from the background, but it can also detect microvessels, predict their correct connectivity and relative position, thus, giving the impression of pseudo-3D vision. The annotation of the original database used for training^[26]^ includes information about spatial location of the vessels with respect to each other: either there is a junction or one vessel pass below another. This information is thus also present in the inferred images and one can distinguish “junctions” and “overlapping vessels”. However, the confidence is based on the quality of the original annotations and thus is not evaluated here. There is no stereoscopic information for spatial location of the vessels, this pseudo-3D is inferred only from one 2D image and has great potential for improving when stereoscopic images of different angles of view are analysed.

**FIGURE 2.**
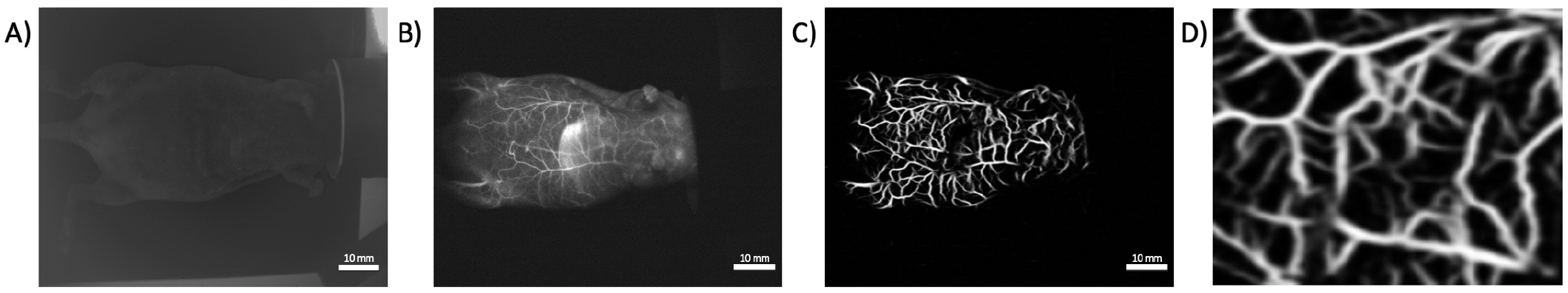
A) SWIR image of a whole body mouse taken at 30 cm working distance before the injection of the contrast agent without long pass filter; B) few seconds after the injection of the contrast agent; C) The resulting prediction of vessels from the image of the whole body mouse in B and D) zoomed image of the segmented blood vessels structure in C.

Figs. 2 A and B shows the vascular network of the ventral side of a whole mouse after the injection of ICG thanks to the detection of its photoluminescence signal above 1500 nm. It should be noted the absence of autofluorescence from the mice before injection in the NIR-IIb region. Vessels of different sizes are detected during the first 4 min after the injection due to the rapid accumulation of ICG in the liver, where it is metabolized by hepatic pathway. Figs. 2 C and D show the segmented vessels from the original SWIR image Fig. 2 B. The vessel structure is well visible with the reduction of signal from the skin and the living tissues. We could see clearlythe anarchic blood network at different depths.

The use a 50 mm lens on the SWIR imaging system allows to reach higher spatial resolution and gain more information on the blood vessels morphology. Fig. 3 top raw depicts the ventral side of the mouse (with consecutive insets) with a high density of blood vessels of different sizes up to micrometer resolution and showing a complex vessels topology. Bottom raw demonstrates the corresponding predicted vessels structure obtained by neural network and provides extremely detailed information not accessible from the raw image by naked eye with junctions and overlaps of the vessels as seen in the insets Figs. 3 B and C.

**FIGURE 3.**
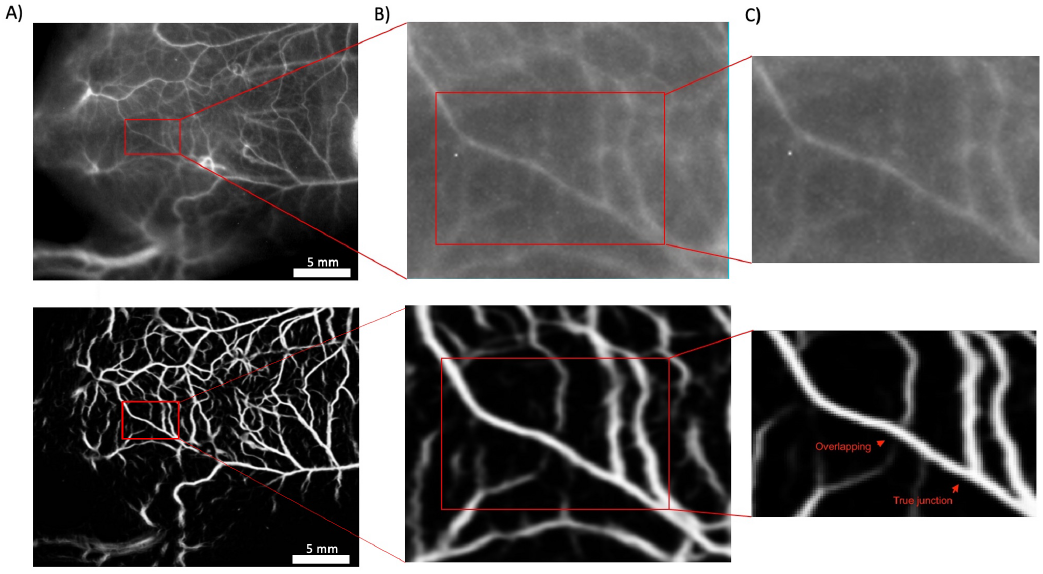
A fragment of the original SWIR image showing complex topology of the blood vessel network with corresponding zoomed insets. It illustrates the predicted blood vessel structure with junctions and cross-sections of the vessels.

Examining the same segmented image Fig. 3 A in inverted colors, Fig. 4 allows to see a complex morphology of blood vessels structure, their relative position, intersection and branching, hence leaving an impression of pseudo-3D shape and depth, as illustrated in the insets of Fig. 4.

**FIGURE 4.**
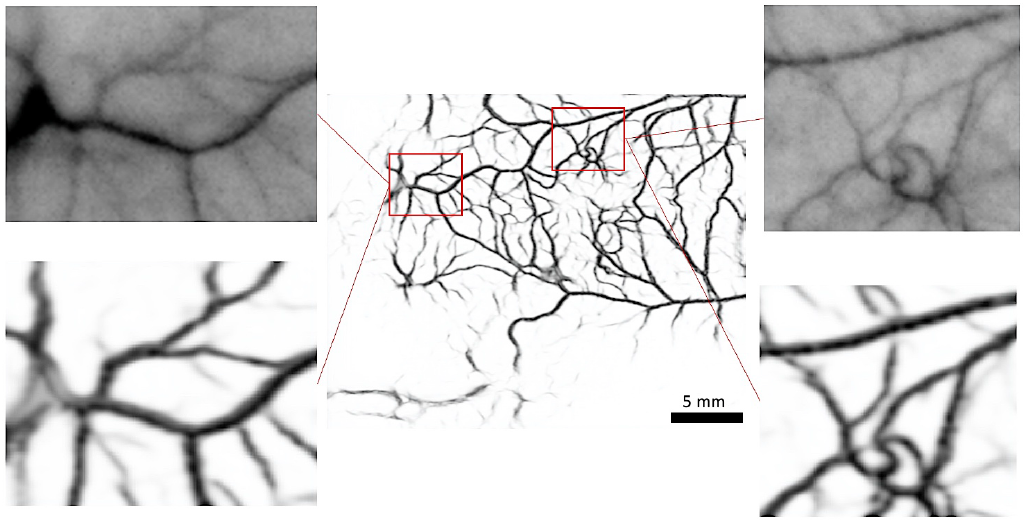
Predicted vessels structure from SWIR image of the whole body of the mouse, Fig. 2 and two fragments in the insets with the detail of the original inverted SWIR images and the corresponding segmented images.

### 2.4 Validation of the vessels morphology

Detection of SWIR signal from vessels below 4 mm depth and keeping a high spatial resolution remains highly challenging. In fact, upper skin is still a main obstacles for SWIR signal, where the light can be scattered and adsorbed. Thus, to confirm microvessels structure predicted by the neural network and inferred from the original SWIR images, one of the direct methods would be the removal of the skin post-mortem and validation of small vessels with high magnification optical camera on *ex vivo* images of the inner skin.

For that purpose, the mice skin flap of 2 to 3 mm thick were soaked in formaldehyde just after the mice were sacrificed. High magnification optical images were taken on the inner side of the flap with an Andor Ikon-M CCD camera at 1 s exposure time with WD 112 mm lens from Leica (zoom x0.8) under white light illumination.

Fig. 5 A corresponds to the optical image of the inner skin of the mouse made with the Andor camera under the white light (neon light which has a broad excitation). A selected area in the center of the inner skin flap (Fig.5 B) was used for inference of the blood vessels with the same IterNet neural network and the same parameters that were used for SWIR images segmentation. The result of the predicted vessels is shown in Fig. 5 C. This image shows main blood vessels and also a number of microvessels. To confirm the existence of these microvessels, three regions of the predicted vessels were selected to compare with separately taken optical images with even higher magnification. The results of this direct comparison are shown in insets with an outline of the same color: red region in Fig. 5 C is compared with an optical image Fig. 5 D; green region Fig. 5 E is compared with optical image Fig. 5 F and blue region Fig. 5 G is compared with Fig. 5 H. It is noteworthy, that optical images that were taken for comparison, are from different images taken on the inner skin sample regions with higher magnification from x1.7 (Fig.5B) to x15.4 (fig.5F), while neural network was applied on Fig. 5 B.

**FIGURE 5.**
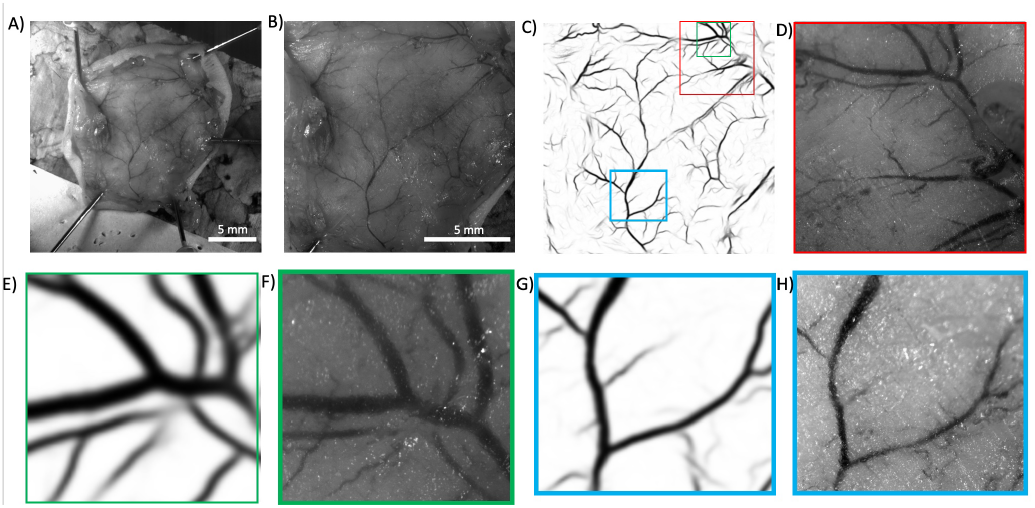
A) An overview of the removed inner skin region; B) Closer view of the removed inner skin region used for inference (zoom x1.7 of image A); C) Predicted vessels structure inferred from B); D) Optical view with higher resolution of the red fragment (zoom x5.6 of image A); E) A green fragment of the predicted vessels; F) The same region in optical image with higher magnification (zoom x15.4 of image A); G) A blue fragment of the predicted vessels; H) The same region in optical image with higher magnification (zoom x8.8 of image A).

This indicates a good confidence to use such neural network for processing SWIR images for inference of blood vessels structure and blood vessel mapping.

## 3 BLOOD VESSELS ANALYSIS

The detection and segmentation of blood vessels are not the only possibilities for the neural network analysis. The propagation of the contrast agent across the blood vessels after the first injection follows non-trivial haemodynamics that can be visualized with an appropriate setup. To demonstrate the capabilities of the method, the initial vascularization of the contrast agent was recorded on the flank of the mouse in the first second after i.v. injection. Very quickly after injection, the constant increase of photoluminescence signal across the vessels in site with high blood vessel density has converted in overexposed spot, where single vessel cannot be visualized. In contrast, the regions, where the contrast agent still has not arrived, are underexposed and there is not enough signal to visualize the vessels, upper raw, Fig. 6 A.

**FIGURE 6.**
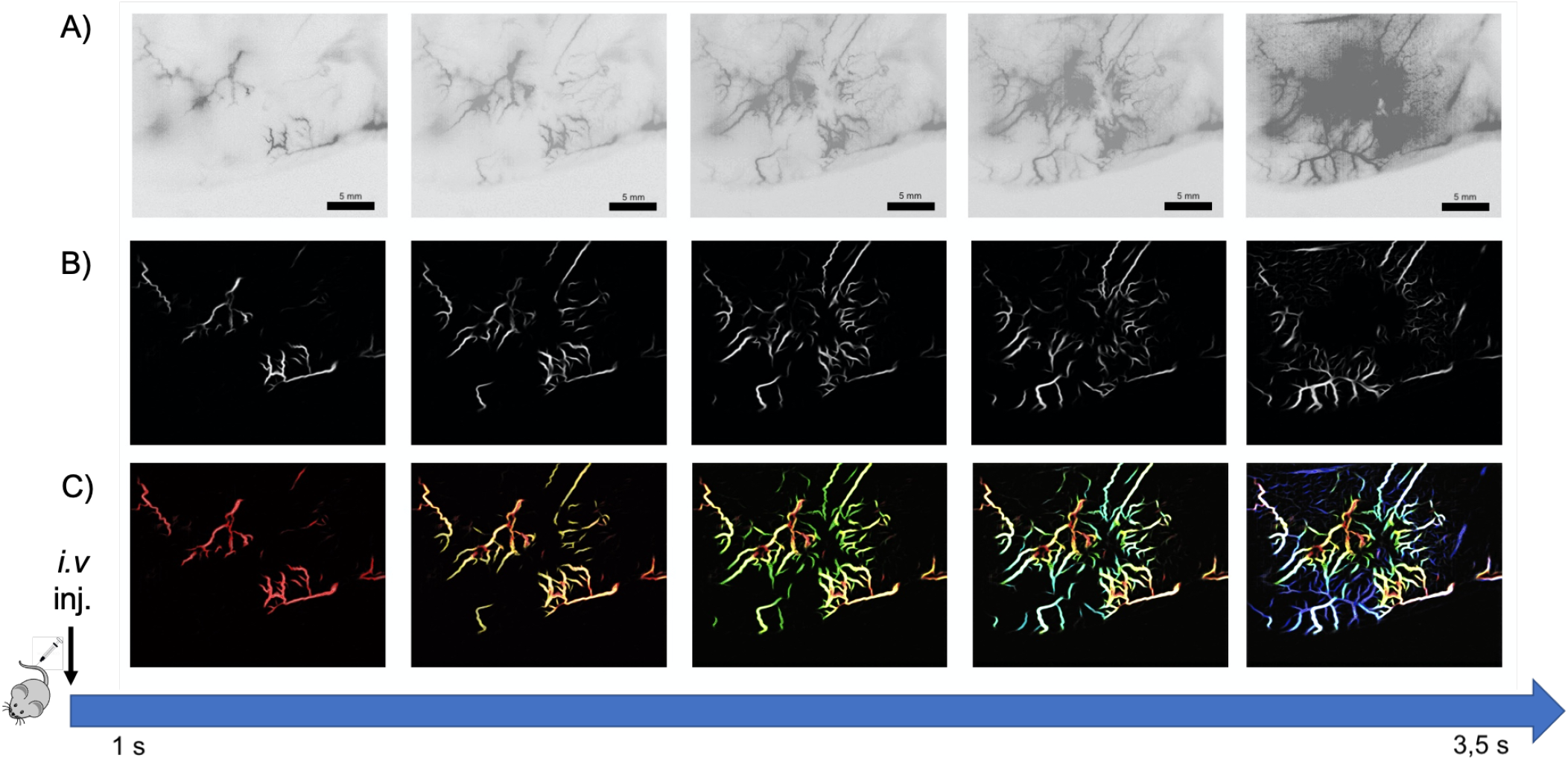
A) Consequent frames (one frame per 500 ms) of the original inverted SWIR image, where the contrast agent propagates through the blood network. B) the corresponding predicted vessels by IterNet network. Vessels in overexposed regions are not detected, thus allowing to follow the blood propagation. C) consequent sum of predicted vessels from individual frames (“SWIR HDR” effect): red corresponds to first frame vessels detection, next frames are superimposed images from previous frames, each with its own color.

The neural network applied to each of the frames does not pick up the whole structure of the vessels, because this information is not present in each image, Fig. 6 B. However, the SWIR images are the sequence of the frames taken with 500 ms time difference of the same region and thus, one can employ a technique, very roughly resembling “High dynamic range” (HDR) image processing in photography, when several images with different exposure are superimposed in one image. SWIR images contain information from different vessels in different time, controlled by the propagation of the contrast agent. Using a different color for each of the frame inference with deep neural network, one can reflect in a combined image not only the whole structure of the blood vessels, but also the colors in the image would reflect the time of the passage of the contrast agent, Fig. 6 C. Thus, combined images allow to follow the kinetics of the circulation of the contrast agent over time during this first 30 s after injection.

Another important feature that can be extracted by neural network is related to the differentiation between arteries and veins. The distinction between veins and arteries is considered critical in angiogenesis and related to the analysis of the couple veins/arteries. This information can be inferred from the shape and the surrounding of the vessels, *e*.*g*. the thick veins are accompanied by thin arteries going parallel to the veins. This information can be learned by the neural network carefully trained on well-annotated examples, where the composition of the vessels is known. This was implemented in the network SeqNet^[31]^, specially trained to distinguish between veins and arteries.

For this purpose we applied SeqNet with provided pretrained weights to SWIR images, that were not optimized and were not included in the training set. Fig. 7 A shows the original inverted SWIR image of the ear mice and Fig. 7 B shows the corresponding segmented image obtained by SeqNet, which segmentation is essentially the same as IterNet. SeqNet network post-processing enables to predict artery and veins based on their size and their distance between each other with high confidence even for images, which are very different from the training set. The resulting distinction in Fig. 7 C-E shows a potential to have a much better differentiation with specially trained networks.

**FIGURE 7.**
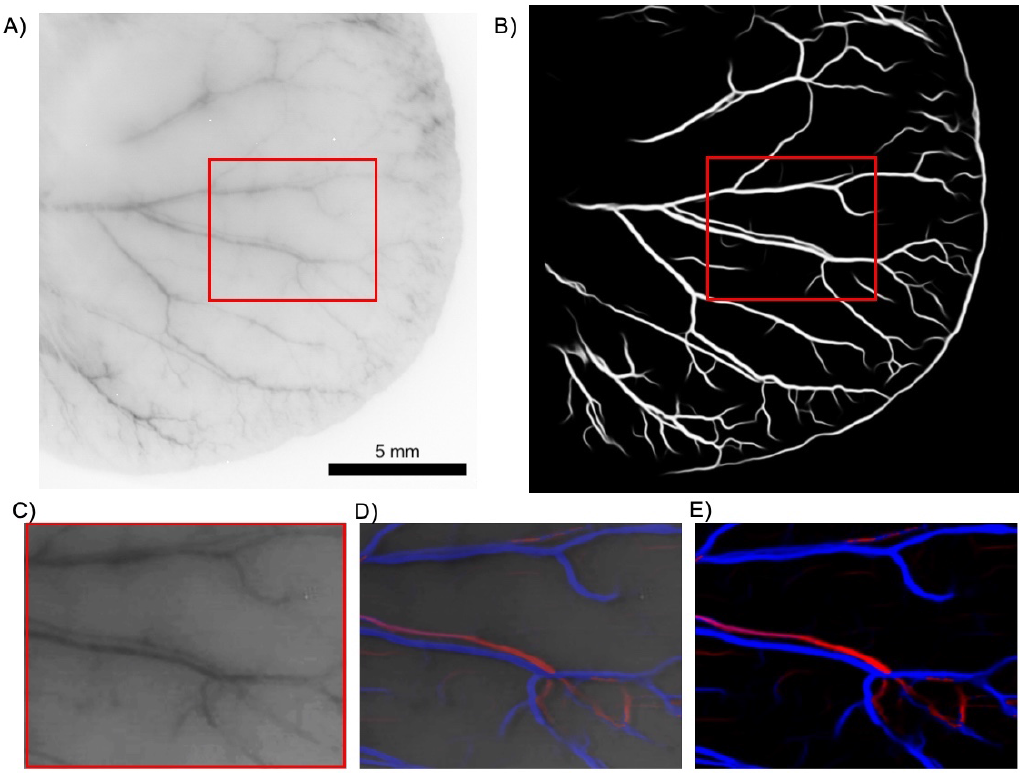
A) Inverted original SWIR image of a mouse ear after ICG injection and B) predicted vessels structure inferred from A. C) Fragment of the original image A. D) Overlay of predicted vessel structure and the original fragment image c. Arteries (red) and veins (blue) predicted by SeqNet^[31]^ neural network; E) Segmented arteries and veins. Predicted arteries (red) and veins (blue) by SeqNet neural network.

The ability to obtain an accurate prediction of the blood vessel network structure by neural networks facilitates further post-analysis of the blood vessels structure tortuosity with more confidence than on the same skeletonization performed on the raw images. This analysis includes extraction of the skeleton of the network and, consequently, deep statistical analysis of network structure: the position and number of branching points, measuring the length of skeleton branches, paths statistics along vessels, density of junctions, vessels, distances between different kinds of structures, comparison of skeletons in order to detect new vessels. As an example, we use a segmentation result of Fig. 1 A as an input for skeleton extraction by open source Python library Skan^[32]^. The result is shown in Fig. 8 and the corresponding video is available in SI. The resulting statistical analysis allows to extract the scaling relations between different types of branches. This, in turn, allows for detailed comparison between sets, for example, it would allow to follow the development of the blood vessels in tumors with comparison against normal tissues.

**FIGURE 8.**
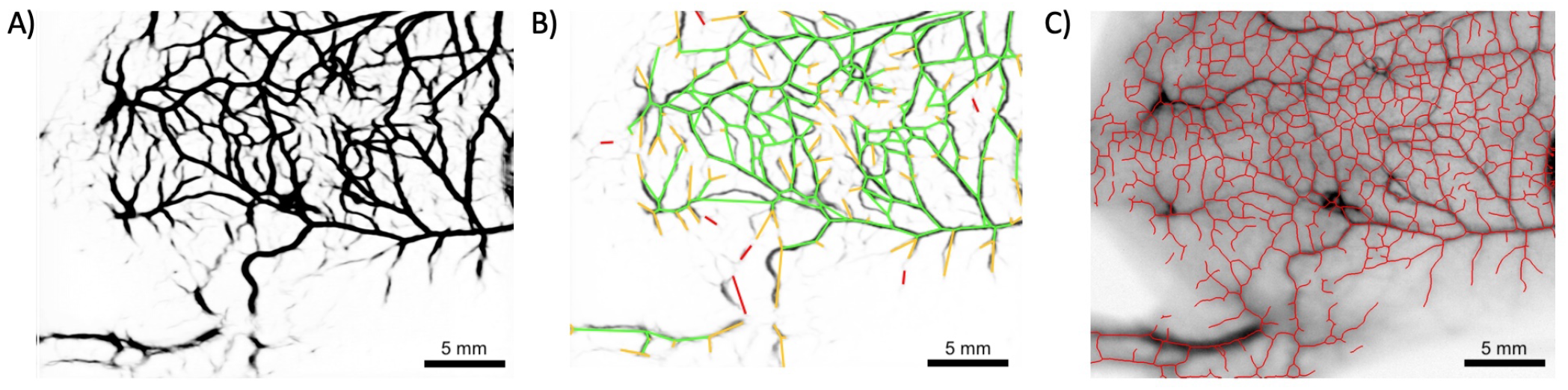
Skeletonization of the predicted blood vessels network. A) Predicted blood vessels from Fig. 1A; B) Euclidean skeleton of the vessels, which connects directly the branching points; C) Vessel network overlapped on the original SWIR image.

## 4 CONCLUSIONS

We demonstrated the potential of deep learning applied to the IR optical imaging in general and in the NIR-IIb (1500-1700 nm) region in particular. This analysis allows to (i) segment vessels from the tissues and noise due to scattering and adsorption of light in tissues; (ii) distinguish vessels overlap and junctions; (iii) to get a morphology and depth estimation with a pseudo-3D shape of blood vessels; (iv) distinguish different hierarchies of vessels types, such as veins and arteries; (v) obtain a information about haemodynamics and kinetics of blood flow; and (vi) perform all types of statistics on vessels junctions, connections, branching using accurate skeletonization of the blood vessel network. Deep learning methods are a clear step to move forward making fast *in vivo* SWIR technique closer to high-resolution optical imaging systems for future biomedical applications.

## Supporting information

segmentation and skeletonization

propagation of blood after injection

## ACKNOWLEDGMENTS

XLG would like to thanks Maxime Henry and Benjamin Musnier for the animal experiments and Plan cancer (C18038CS) for their financial support. The Optimal imaging platform is supported by France Life Imaging (French program “Investissement d’Avenir” grant; “Infrastructure d’avenir en Biologie Santé”, ANR-11-INBS-0006) and the IBISA French consortium “Infrastructures en Biologie Santé et Agronomie”.

## ETHICS STATEMENT

Animal experimentations were performed in accordance with the institutional guidelines of the European Community (EU Directive 2010/63/EU) for the use of experimental animals and received the approval of the local ethics committee (Cometh38 Grenoble, France) and the French Ministry of Higher Education and Research under the reference: APAFIS#21916-2019082710189095 v4.

## AUTHORS CONTRIBUTIONS

Computational analysis and methodology: Vladimir Baulin. Analysis and interpretation of data: Vladimir Baulin and Xavier Le Guevel. Drafting of the manuscript: Vladimir Baulin and Xavier Le Guevel. Animal experiments and imaging: Xavier Le Guevel, Yves Usson. Critical revision of the manuscript for important intellectual content: all authors. Obtained funding: Xavier Le Guevel.

## DATA AVAILABILITY STATEMENT

The data that support the findings of this study are available from the corresponding author upon reasonable request.”

## SUPPORTING INFORMATION

Analysis of the SWIR video with deep learning and skeletonization algorithms related to Fig. 8 and the segmented video of blood propagation through vessels, related to Fig. 6.

## Graphical Abstract Figure

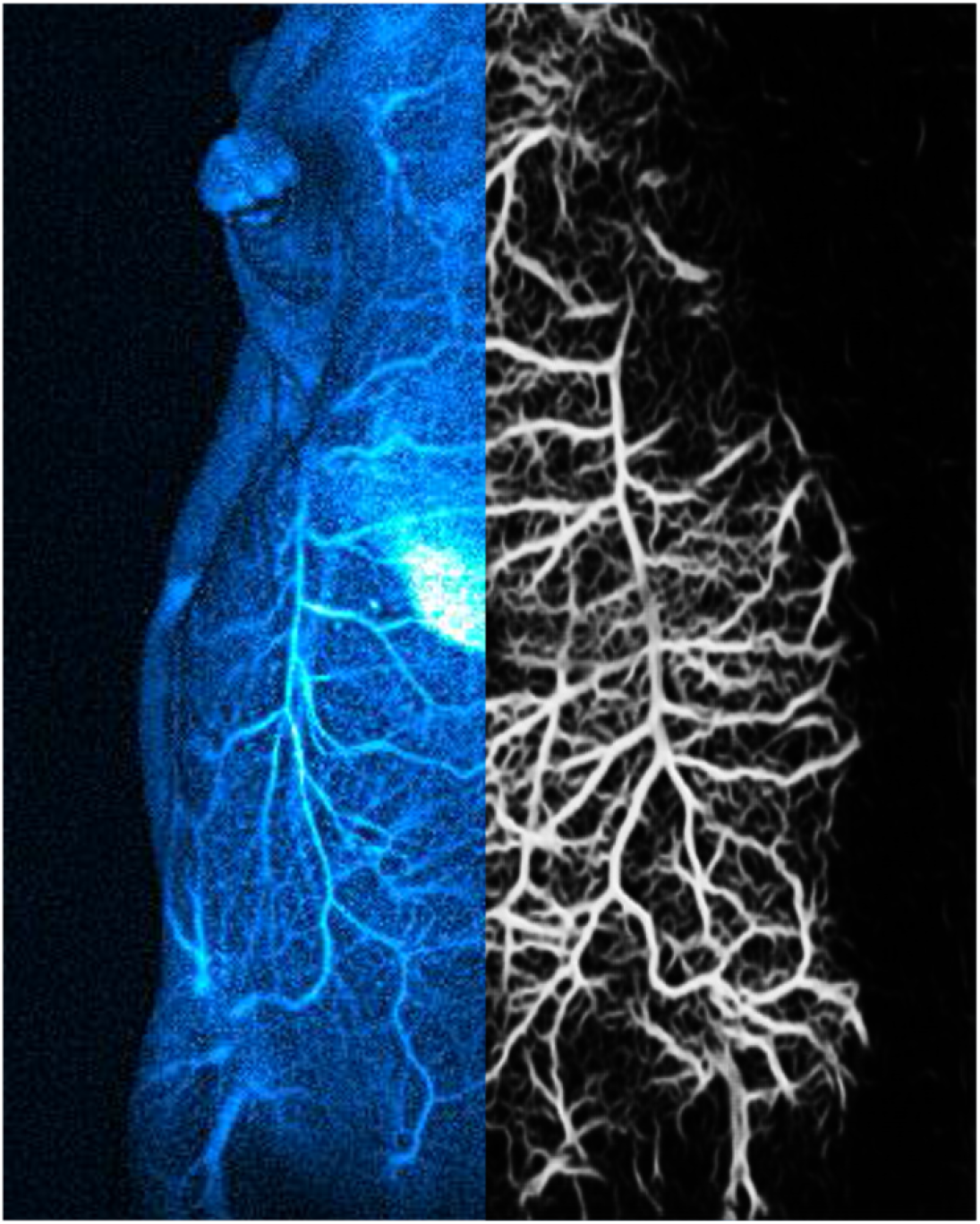
Deep learning methods provide a fast approach to move noninvasive infrared in vivo imaging to high resolution imaging techniques with full analyses of the vascular network

